# Requirements for swarming ability by lateral flagella on an agar surface in marine *Vibrio* cells

**DOI:** 10.64898/2026.08.08.743661

**Authors:** Michio Homma, Takehiko Mima, Hajime Nakatani, Seiji Kojima

## Abstract

The marine bacterium *Vibrio alginolyticus* and the food poisoning bacterium *V. parahaemolyticus* possess two types of flagella in one cell: proton-driven lateral flagella (Laf) extending from the periphery of the cell body, and sodium ion-driven polar flagella (Pof) extending from a cell pole. For swimming in seawater they use polar flagella, suppressing the expression of lateral flagella. When they attach to the body surface of fish or intestinal tract, lateral flagella are induced, allowing it to crawl along the surface or through mucus. The dynamometer hypothesis, which proposes that polar flagella sense rotation and control the expression of lateral flagellar genes, has been widely accepted. However, how rotation is sensed and how expression is controlled remains unclear. Although swarming has recently been analyzed by physical, biological, or biochemical perspectives, it remains unclear how this motility is controlled, or which substances and conditions are necessary for swarming ability. In this study, we discovered that adding gelatin to agar medium promotes swarming on the agar surface by the lateral flagella of *Vibrio*. Our data suggested that surfactants or viscous polysaccharides secreted extracellularly are important for promoting swarming on the agar surface and we identified that swarming is likely to be driven by S (social)-motility, in which bacteria move by interacting with each other, and A (adventure)-motility, in which bacteria move by interacting with the agar surface. Our study provides clues that help clarify the mechanism of bacterial swarming

**IMPORTANCE:** We discovered that adding gelatin to hard agar medium promoted swarming on agar surfaces by the lateral flagella of *Vibrio* cells. The surfactants or viscous polysaccharides secreted extracellularly seem to be important for swarming ability on agar surfaces. We proposed that the swarming is thought to occur through S(social)-motility, where cells move by interacting with cell bodies each other, and A(adventure)-motility, where cells move by interacting with the agar surface and cell body. The present study should provide the clues to clarify the mechanism of bacterial swarming and how to move in a viscous environment.

## INTRODUCTION

Bacteria are known to swim in liquid media, but the ability to swarm on solid media or an agar plate was identified first in *Proteus* bacteria [1, 2]. Subsequently surface swarming was found to be widespread in flagellated bacteria [3, 4]. Bacterial movement on surfaces such as agar can occur by multiple mechanisms, including the extension and retraction of specialized pili (“twitching”) [5, 6], the movement of membrane-bound adhesins along the cell surface (“gliding”) [7, 8], a detergent-assisted but primarily passive growth mechanism (“sliding”) [9], or swimming in a thin layer of fluid at the surface (“swarming”) [10, 11]. Sliding and swarming on agar surfaces have been observed in a variety of genera, including *Proteus* [1], *Vibrio* [12, 13], and *Bacillus* [14], as well as the Gram-negative model species *Salmonella enterica* and *Escherichia coli* [15]. Although the community or colony migration of *S. enterica* and *E. coli* has been extensively studied, the mechanisms by which they crawl on an agar surface are not completely understood [16–18]. *S. enterica* and *E. coli* require relatively low agar concentrations (approximately 0.5%) for migration on the agar surface. In contrast, *Proteus mirabilis* can migrate on harder surfaces (>2% agar) [19]. While swarming of *B. subtilis*, *Staphylococcus aureus*, and *Pseudomonas aeruginosa* depends on cell-produced surfactants, *E. coli* and *S. enterica* swarming are thought to rely on cell-derived osmotic agents that draw water from the medium and form a wet zone around the edge of the expanding colony [20, 21]. However, the osmotic agents have not been identified. Flagella-deficient and motility-deficient strains do not swarm, indicating that cell migration within this fluid layer depends on active propulsion by flagella.

Many bacteria can move using organelles called flagella, which are driven by sodium ion or proton ion flow [22, 23]. Flagella consist of a filament acting as a screw, a motor forming the engine and a hook that connects the motor and screw. The motor can rotate in a counterclockwise (CCW) or clockwise (CW) direction [24, 25]. In peritrichous bacteria such as *E. coli*, CCW rotation results in bundling of flagellar filaments that pushes the cell body for smooth swimming, while CW rotation causes the flagella to pull the cell body, inducing tumbling movement that changes the swimming direction. By controlling the rotational direction of the motor, the cell moves towards favorable environments or can escape from unfavorable environments. In bacteria like *Vibrio cholerae*, which possess a single flagellum at the pole, CW rotation of the motor pulls the cell body, causing it to move backwards.

The flagellar motor consists of a rotor and surrounding stator subunits [26, 27]. The stators are ion channels that convert the electrochemical potential energy of protons or sodium ions across the cell membrane into kinetic energy. The rotor comprises an MS ring embedded in the cell membrane and a cytoplasmic C-ring attached beneath the MS-ring. The stator is composed of two types of membrane proteins, the A subunit and the B subunit [28]. For example, MotA and MotB form the proton-driven stator in *E. coli*, PomA and PomB form the sodium-driven stator in *Vibrio*, or MotP and MotS form the sodium-driven stator in *Bacillus*. The A subunit (MotA/PomA/MotP) comprises four transmembrane helices and a large cytoplasmic domain which contains highly conserved charged residues essential for torque generation and interacts with the C-ring component protein FliG to drive the rotor rotation. The B subunit (MotB/PomB/MotS) consists of a single transmembrane helix and a C-terminal OmpA-like domain. This transmembrane helix contains conserved aspartic acid residues essential for the ion channel activity of the stator. The OmpA-like domain binds to the peptidoglycan layer or T-ring, anchoring the stator complexes around the rotor [29].

The marine *Vibrio* (*V. alginolyticus*) and the food poisoning bacterium (*V. parahaemolyticus*) both possess two types of flagella in one cell: proton-driven lateral flagella located around the periphery of the cell and sodium ion-driven polar flagella located at the cell poles [30, 31]. When swimming in seawater, they use polar flagella and the lateral flagella are suppressed. While attached to the surface of a fish or the intestinal tract, lateral flagella are induced, allowing them to crawl along the surface or through mucus [32]. Expression of the lateral flagella is also induced by Amiloride derivatives, sodium channel inhibitors, that inhibit sodium-driven polar flagellar motility [33]. These observations led to the proposal of the widely accepted dynamometer hypothesis, which states that flagella can sense rotation and control flagellar gene expression [34, 35]. However, how rotation is sensed and how expression is regulated remains unclear. Motility via lateral flagella is possible on the agar surface of hard agar plates (1.25%), but not on the agar surfaces of 2.5% or higher of agar [10, 36]. This phenomenon of crawling on the agar surface has also not been fully explained.

FliL has been identified as a protein that assists stator function [37]. *V. alginolyticus* has two types of FliL for polar and lateral flagella. Mutants of FliL (^pof^*fliL* and ^laf^*fliL*) exhibit reduced polar flagellar motility at high viscosity and defective in lateral flagellar motility on hard agar plate, respectively [38]. Therefore, FliL is closely involved in the swarming ability of lateral flagella. FliL is a single-pass transmembrane protein reported to be located near the stator and basal body, interacting with them [39, 40]. It has also been shown that FliL is important for the efficient incorporation of the stator into the motor [39]. A possible model has been proposed that FliL interacts with the stator to form a stator-FliL complex surrounding the stator [38]. Structural analysis revealed that FliL has a stomatin/prohibitin/flotillin/HflK/C (SPFH) domain, which is widely conserved from mammals to bacteria. The SPFH domain is also known to be involved in mechanosensing and ion channel regulation [41, 42].

Surface swarming by flagella has recently been analyzed from a physics point of view [43, 44]. However, from the biological or biochemical perspective, it is still unclear how the swarming motility is regulated and what materials or conditions are necessary for the swarming ability. In this study, we found that the addition of gelatin in hard agar plates promotes swarming on the surface of agar by marine *Vibrio* lateral flagella. We suggest that surfactants or viscous polysaccharides secreted outside the bacterial cell are important for the swarming motility on the agar surface.

## RESULTS

### Swarming ability induced by gelatin

The motility of *Vibrio alginolyticu*s has been previously analyzed using polar and lateral flagellar mutants [31]. In that report, the motility on agar plate was examined using heart infusion broth (Difco) containing 1.5% agar. Moreover, it was shown that lateral flagellar deficient mutants prevented the motility on 1.5 % agar, even polar flagella are present. Lateral flagella were also shown to be capable of motility in highly viscous environments containing polyvinylpyrrolidone (PVP), a water-soluble polymer that confers buffer or medium with high viscosity [32]. Here, we discovered the medium conditions required for swarming motility on 1.25% agar for the first time with the reproducible swarming profiles (Fig. 1). Overnight cultures of the wild-type (wt) strain (138-2), the polar flagella (*pof*)-deficient strain (YM19), and the polar flagella and lateral *fliL* (*pof* ^laf^*fliL*)-deficient strain (YM19Δ*fliL*, named NMB342) were spotted onto 1.25% agar VNG plate and incubated at 30°C for 6h. The swarming ring formation by bacterial spreading was limited on the media containing only casein hydrolysate as a nutrient source (Fig. 1C), but was significantly greater on media containing bovine gelatin (Fig. 1A). The ^laf^*fliL* mutant (shown as c in Fig. 1) showed very limited spreading. After extended incubation, even in the agar plate without gelatin, the bacterial cells except the ^laf^*fliL* mutant were able to spread on the 1.25% agar (Fig. 1D). This suggested that the addition of gelatin induced the release of products that enabled spreading of the swarming ring on the agar surface. The induction of swarming ability also occurred with the addition of chitin oligosaccharides and collagen peptides, the hydrolyzed products of bovine or fish gelatin (Supplementary Fig. S1). Furthermore, adding PVP (final concentration of 5%) to the 1.25% agar media enabled bacterial spreading on the agar surface, similar to profiles by adding gelatin (Fig. 2). The ^laf^*fliL* mutant did not induce swarming by PVP. We hypothesized that gelatin may induce the secretion of a highly viscous substance such as PVP outside the cell.

**FIG 1.**
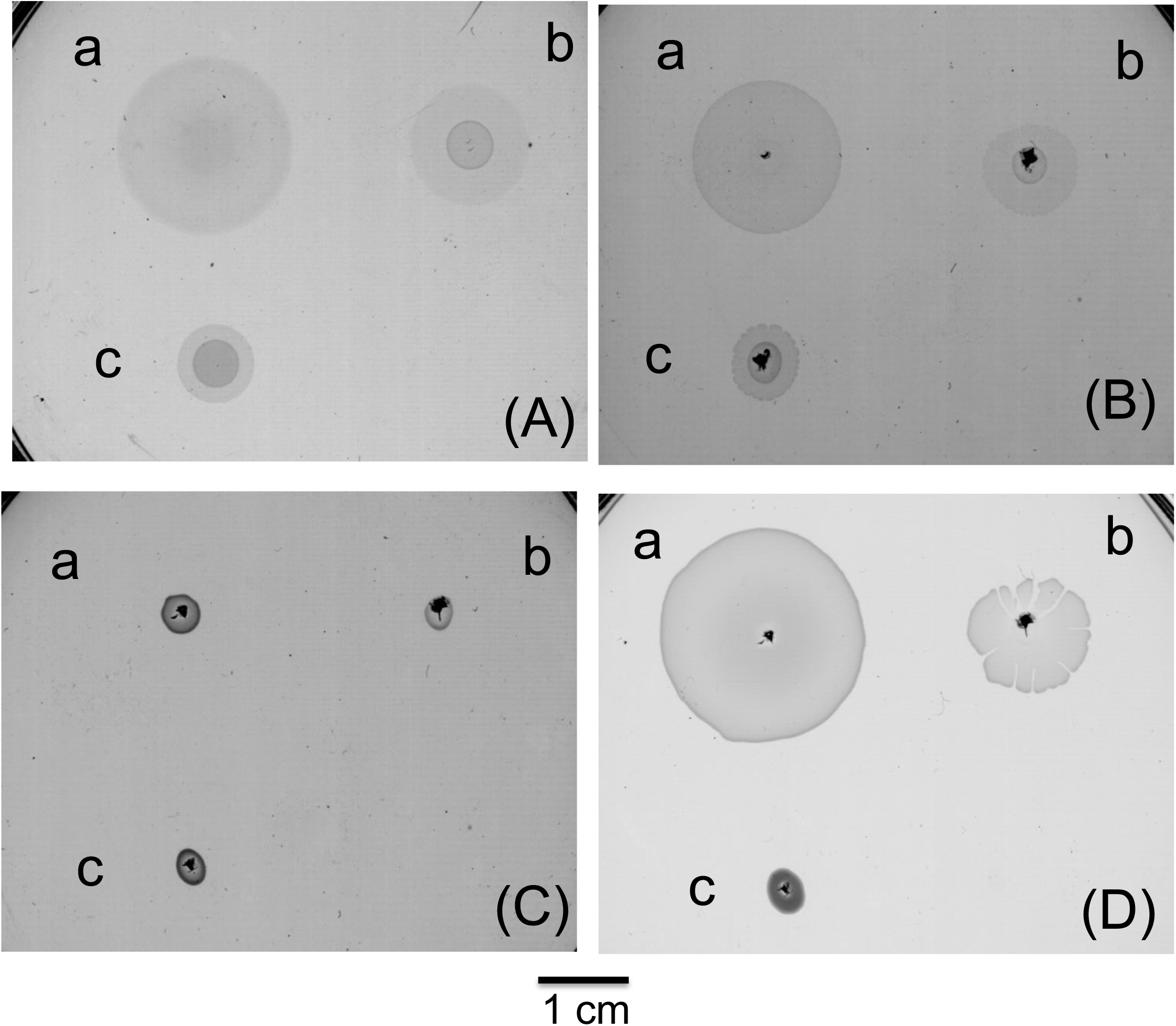
The swarming by gelatin addition. 2 μL of the overnight culture (a:138-2, b:YM19, c:YM19^laf^*fliL*) was applied to the 1.25% agar-VNG plate containing 1% gelatin (A), 1% collagen peptides (B), 1% casamino acid (C) and incubated at 30°C. The plates were scanned with a photo scanner after 6 hours (A-C) and (C) was further incubated for 10 hours (D).

**FIG 2.**
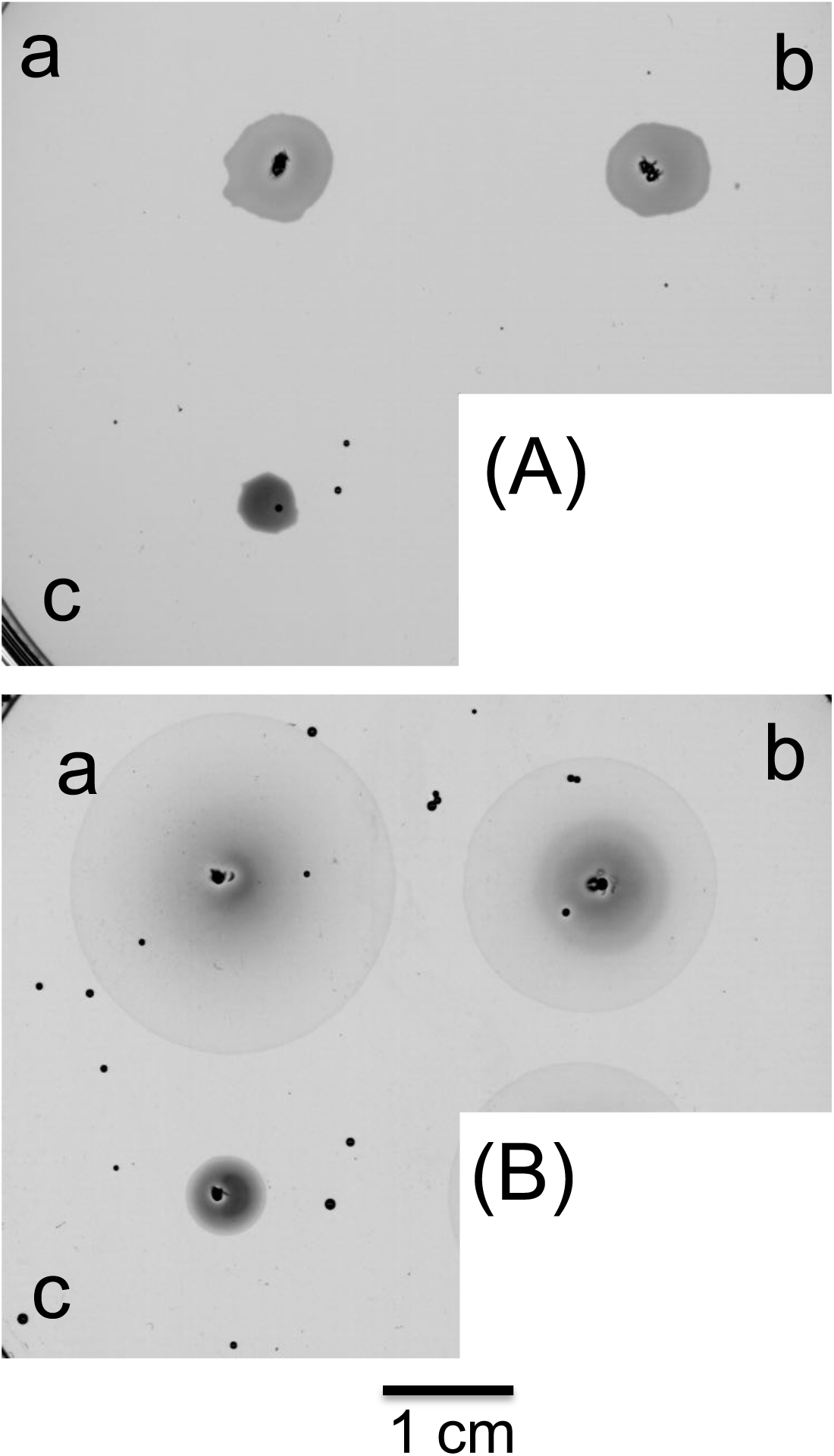
The swarming by PVP addition. 2 μL of the overnight culture (a:138-2, b:YM19, c:YM19^laf^*fliL*) was put on the 1.25% agar-VNG plate (A), or containing 5% PVP (B). The plates were scanned with a photo scanner after 6 hours.

Next, we observed the flagellar formation of swarming cells on the agar plates. After overnight cultures were spotted onto the 1.25% agar VPG plate and incubated at 30°C for 6h, the edges of the colonies were observed using an electron microscope (Supplementary Fig. S2). In the wild-type strain (138-2), numerous lateral flagella were observed around the cell periphery, whereas polar flagellum was not observed or distinguished in the present condition. In the *pof* deficient strain (YM19), numerous lateral flagella were observed around the cell periphery as in wild-type cells, but the polar flagellum was not observed. In the *pof* ^laf^*fliL*-deficient strain (NMB342), numerous lateral flagella were observed around the cell periphery, although the number of lateral flagella seems to be slightly lower than in the *pof*-deficient parent strain YM19.

We directly observed cell motility on the agar. After overnight cultures were spotted onto 1.25% agar and incubated at 30°C for 6h, the edges of the colonies were observed cells using a digital microscope (Fig. 3, Supplementary Fig. S3, movies that demonstrate motility at the edge are included). On the gelatin-containing agar medium, the wild-type and *pof*-deficient strains, whose cells were elongated and in a mostly single layer, moved through wriggling motion (Fig. S3a and S3b). The *pof* ^laf^*fliL*-deficient strain moved very slowly and the cells were multi-layered due to their inability to move outward (Fig. S3c). Areas other than the edge of the swarming ring of the *pof*-deficient strain on 1.25% agar medium were also observed using digital microscope video (Supplementary Fig. S4). The size and motility of the cells are different among the places of ring. Other than the edge regions, bacterial cells were multi-layered, elongated and moving everywhere in all locations. Cells of the inner region appear less elongated because the cells had been divided.

**FIG 3.**
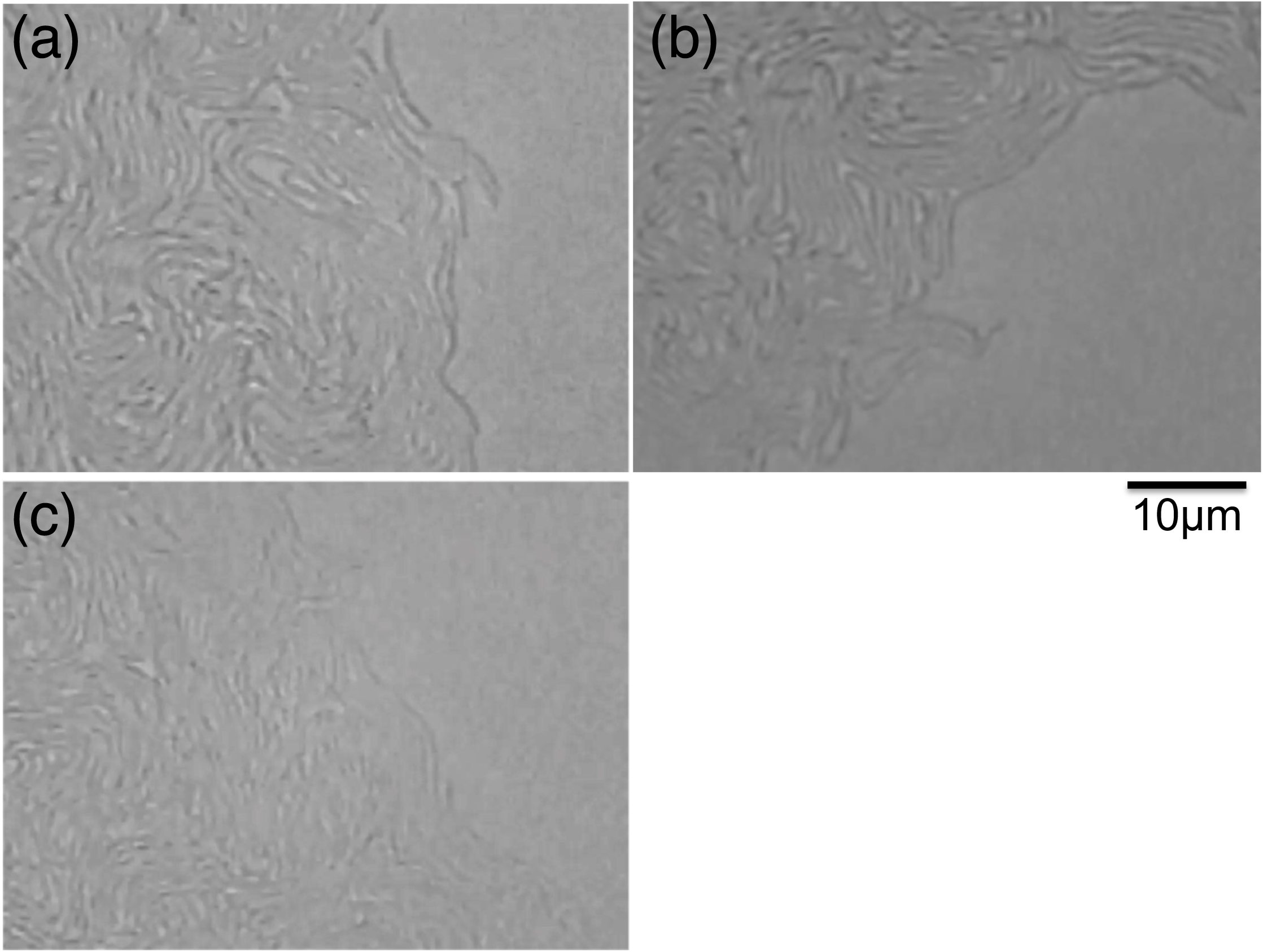
Swarming profiles of cells on an agar plate. 2 μL of the overnight culture (a:138-2, b:YM19, c:YM19^laf^*fliL*) was inoculated and incubated at 30°C for 6 hours on the 1.25% agar-VNG plate containing 1% gelatin. The edges of swarming rings (for 138-2 and YM19) or colony (YM19^laf^*fliL*) were observed by digital microscope (see also Fig. S3).

### Swimming ability in the presence of PVP

It has previously been shown that the viscosity of the medium significantly affects the swimming motility of *V. alginolyticus* cells [32]. Here, in wild-type cells, which possesses both polar and lateral flagella, only the polar flagella were constitutively expressed in low-viscosity media, resulting in forward and backward swimming with a characteristic phenotype by polar flagella (Fig. 4a, Supplementary Fig. S5a). On the other hand, in high-viscosity media containing PVP, the lateral flagella were expressed and the bacteria swam smoothly (Figs. 4d, Supplementary Fig. S5d). However, in the *pof*-deficient strain (YM19), the lateral flagella were expressed even in low-viscosity media, resulting in motility with reduced frequency of directional changes (Fig. 4b, Supplementary Fig. S5b). By contrast, the *pof* ^laf^*fliL*-deficient strain (NMB342) was almost completely non-motile in low-viscosity media, but showed enhanced motility in high-viscosity media (Figs. 4c and f, Supplementary Fig. S5c and f). This evidence may contradict the previous reports suggesting that FliL is essential for resisting the stress by high viscosity.

**FIG 4.**
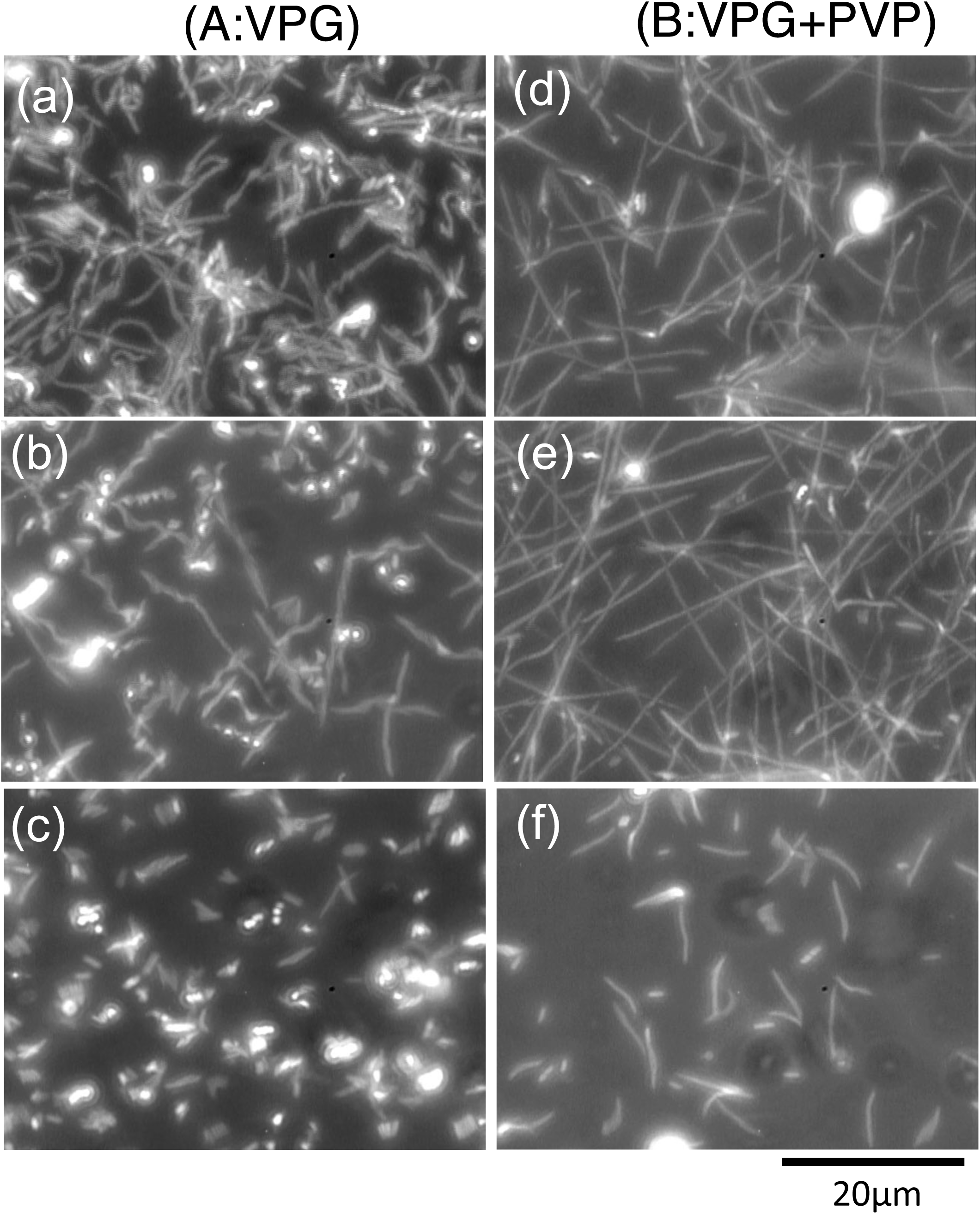
Swimming motility in a culture with or without PVP. 15 μL of the overnight culture (a,d:138-2, b,e:YM19, c,f:YM19^laf^*fliL*) was inoculated into 1mL of VPG (A) or VPG containing 5% PVP (B) and incubated at 30°C for 3 hours. The cells were observed with a cover glass by dark field microscopy and recorded by CCD camera. The 30 images (1 sec) were integrated using a motion analysis software.

### Swarming and swimming ability of *fliL* mutant

We examined the function of ^laf^*fliL* using the *pof* mutant cells (Fig. 5). On 1.25% agar, the swarming ring of the ^laf^*fliL* mutant did not expand, but the mutant cells with the ^laf^*fliL* gene introduced had an expanded swarming ring. On 0.3% agar, the swimming ring of the ^laf^*fliL* mutant was slightly smaller than that of the mutant cells with the ^laf^*fliL* gene, although they exhibited good motility. This confirmed that the ^laf^*fliL* mutation plays a major role in the swarming ability on the agar surface. Digital microscope video observation of the edge of the swimming ring revealed that the ^laf^*fliL* mutant and the ^laf^*fliL* mutant complemented by the ^laf^*fliL* gene gave the almost identical swimming motility in soft agar, confirming that the ^laf^*fliL* gene does not contribute the swimming ability (Supplementary Fig. S6).

**FIG 5.**
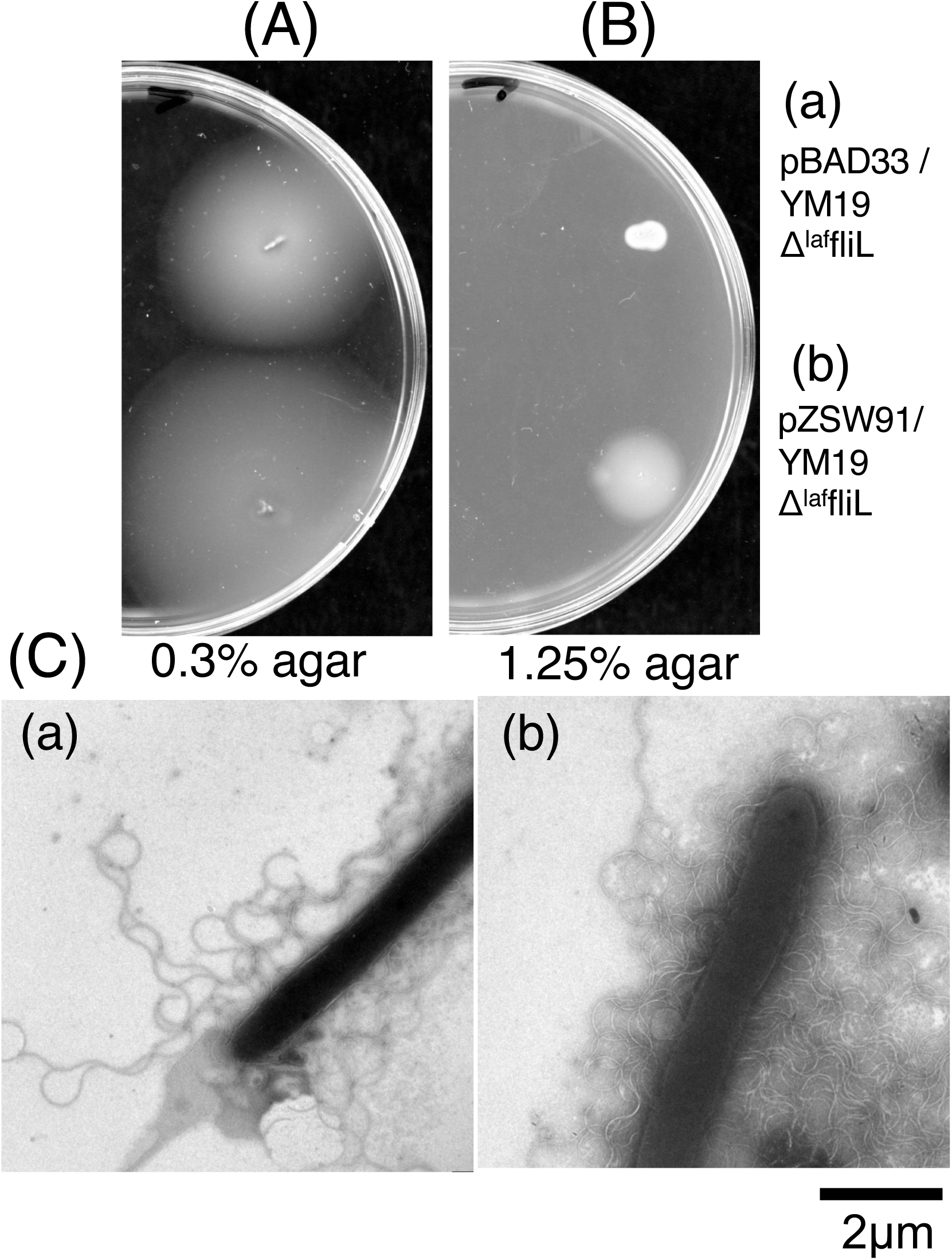
Effect of *fliL* gene for motility. 2 μL of the overnight culture of YM19Δ^laf^*fliL* harboring pBAD33 (a) and pZSW91 (b) was applied to the 0.3 % agar-VPG plate (A) or 1.25% agar-VPG plate (B), containing 2.5μg/mL Cm and 0.1% Arabinose and incubated at 30. The plates were scanned with a photo scanner after 24 hours. The edges of the colonies from (B) were picked up with a toothpick, suspended in 0.5% PTA, and observed under an electron microscope (C).

### Extracellular materials of the swarming cells

The cells grown on 1.25% agar medium were harvested and homogenized to obtain extracellular materials. The separated materials were detected by SDS-PAGE, followed by CBB staining and silver staining (Fig. 6). The most abundant protein in the extracellular materials seems to be flagellin (LafA) indicated by open arrow in Fig. 6B. The staining profiles are similar either in the presence or absence of gelatin though the high molecular region (indicated by closed arrows) stained by silver in the presence of gelatin is denser than that in the absence of gelatin. This might indicate that the size of extracellular polysaccharides is increased by addition of gelatin.

**FIG 6.**
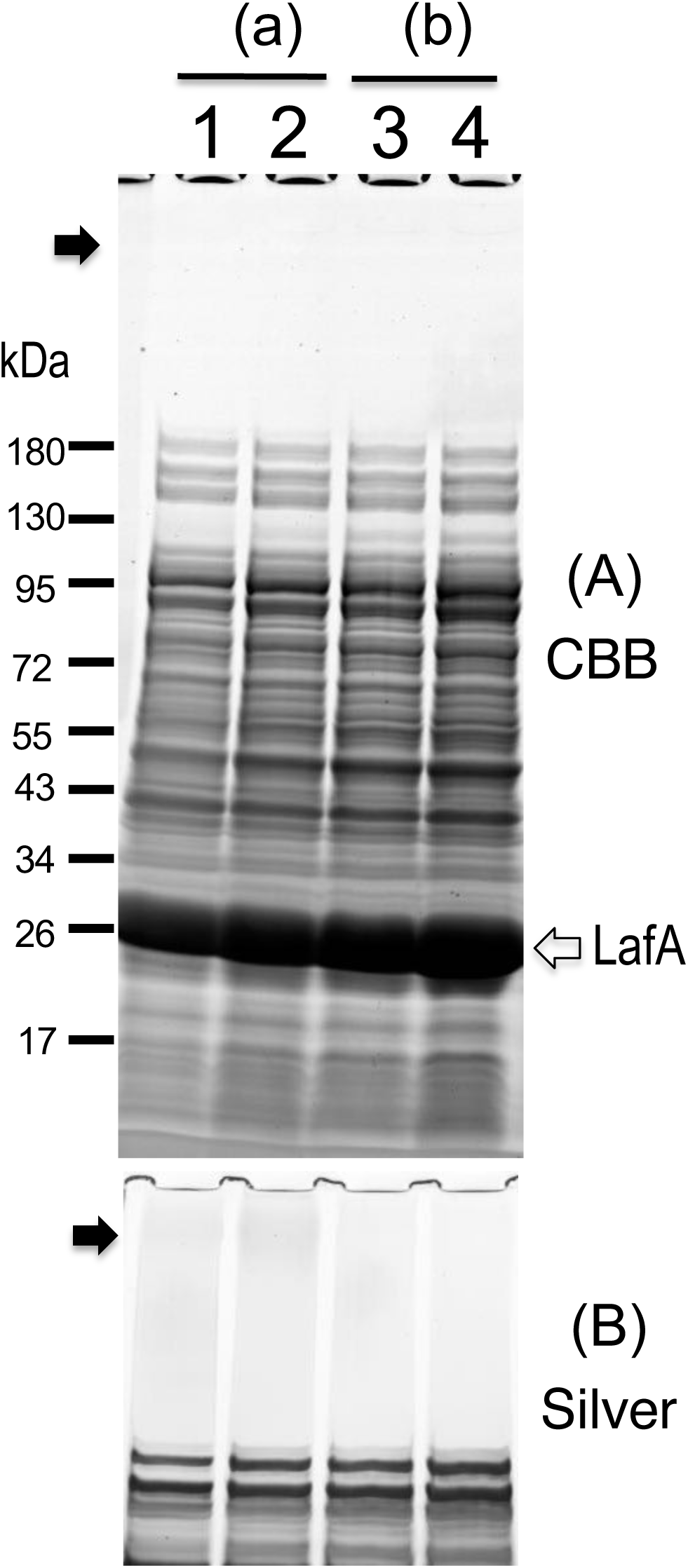
SDS-PAGE profiles of cell surface materials. 50 μL of overnight culture YM19 (1, 2) and YM19Δ*fliL* (3, 4) were inoculated onto 1.25% agar-VNG plates (+none:1, 3) and (+1% Gel:2, 4), and incubated at 30°C. After 6 hours, the cells were scraped and suspend in 1 mL of 20TN200. The cells were stirred using a homogenizer, and removed by low-speed centrifugation. The supernatant was collected as the samples and was separated by SDS-PAGE, stained with Coomassie Brilliant Blue (A: CBB) and silver (B: Silver), and photographed.

## DISCUSSION

Previous studies have shown that swarming on agar in various bacteria is significantly affected by various factors such as temperature, salt concentration, and agar concentration. In this study, we found that the swarming ability on agar surface by the lateral flagella of *V. alginolyticus* is significantly affected by the addition of gelatin. The swarming was also promoted by the addition of gelatin degradation products (collagen peptides), or a chitin degradation product, although the swarming profiles are slightly different. Swarming edge was distorted when fish collagen peptides are included in the plate (Fig. S1c), and the lack of ^laf^*fliL* affected swarming more severely when chitin is included in the plate (Fig, S1a). We do not know the actual reason for the difference, however, we speculate that the additives on the plate would affect somewhat differently the expressions of flagellar genes or of genes to support the swarming, such as the polysaccharide synthesis genes. Furthermore, the addition of the viscous substance PVP also promoted swarming without gelatin. These findings imply that the secretion of viscous substances from the bacterial cell is necessary for the swarming promotion on agar surface. Factors involved in biofilm formation are thought to influence the production of extracellular materials and affect the swarming or swimming by flagella [45, 46]. Surfactant secretion could also be a factor promoting swarming. It has been reported that the addition of Tween-80 to agar medium promotes the formation of swarming rings [47]. Comparison of extracellular substances from cells grown between in the absence and in the presence gelatin, swarming-promoting factor revealed almost no significant difference in proteins stained by CBB or polysaccharides stained by silver staining though polysaccharides of high-molecular weight seem to be increased a little. We plan to conduct a more detailed analysis for the extracellular substances in future.

From the observations of the movement of colony cells, there appear to be two ways in which the propulsive force is generated: (i) interactions between the cell surface and the agar surface, and (ii) surface interactions between cells themselves (Fig. 7). We can imagine that flagellar filament is the tire of car and the agar surface is the road surface. These two kinds of movements phenotypically resemble motility of *Myxococcus*. When a single cell or the small group of *Myxococcus* cells move, they exhibit A (Adventure)-motility, whereas when they move as a large group, they show different motility, called S (Social)-motility [48, 49]. It has been proposed that the swarming motility of *S. enterica* and *E. coli* is driven by flagella swarming through the water on the agar surface [20]. In our observations in this study, vigorous cell movements by the interaction with elongated cells did not promote the ring formation or the swarming on agar plates. It is likely that interactions on the cell surface are responsible for the motility. Flagella, which are locomotive machines, are located in the outermost part of the cell, and we could imagine that flagella interact with each other to drive motility. Interactions between the agar surface and the cell surface or flagella seem to promote rapid spreading on agar surface (A-motility), forming a single cell layer that spreads outward from the place where the cells were spotted. Depending on the conditions of the agar surface and/or of the cell surface, the driving force is not generated between the cell surface and the agar surface, thus A-motility does not occur, and it is possible to spread the swarming ring formed by movements of the cell-cell interactions via S-motility. The expansion of the cell ring due to S-motility is slower than that due to A-motility, and the cells spread in multiple layers.

**FIG 7.**
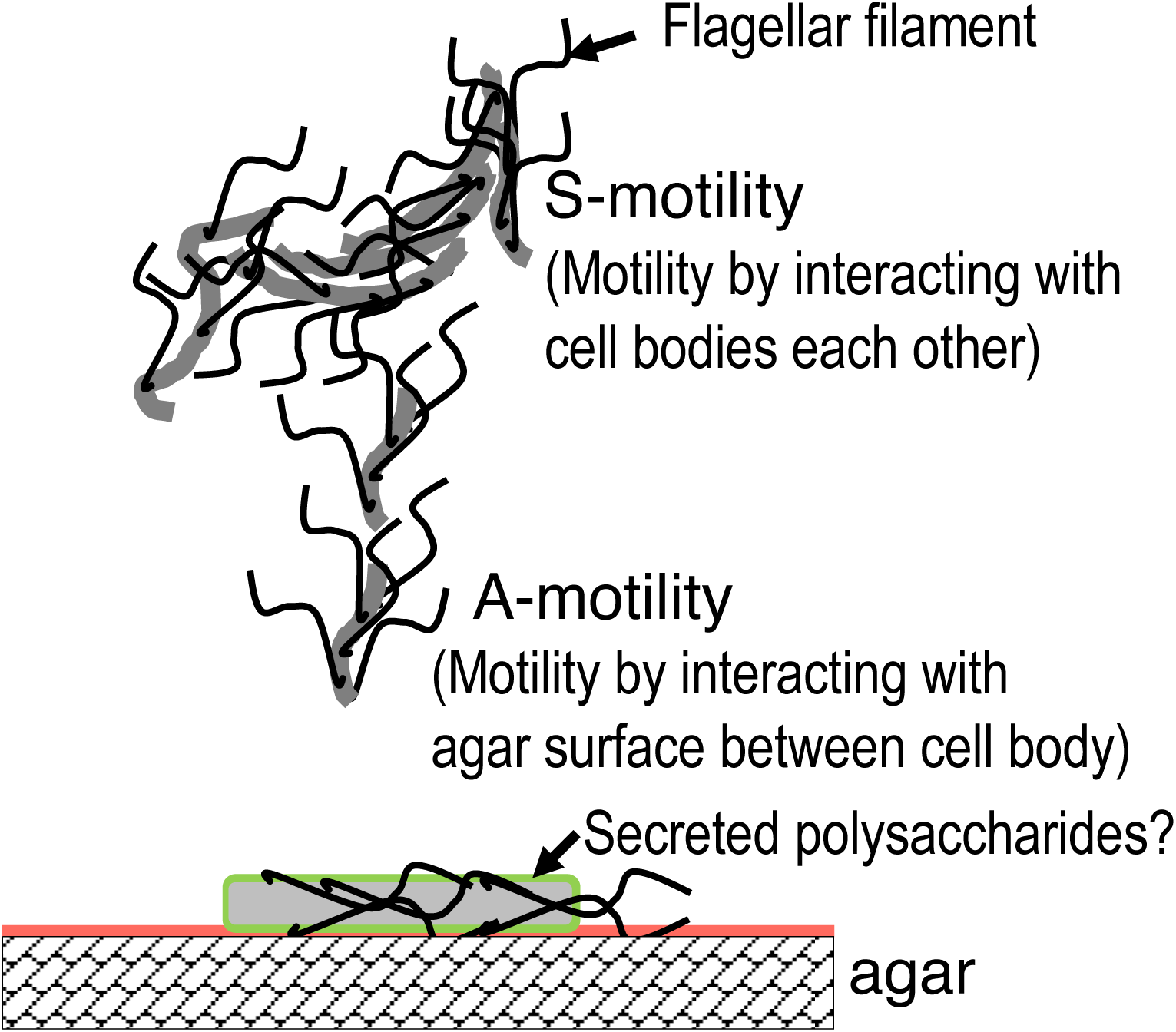
Model of cell motility by lateral flagella on agar. The water layer on agar and the cell surface layer are shown as orange line and the green line, respectively. The cells and flagellar filaments are shown by gray lines and the black lines, respectively.

It had been shown that the expression of lateral flagella is required for cell migration on plates [50]. In the wild-type strain (138-2), lateral flagellar expression is suppressed in liquid medium, and cell movement is achieved by constitutively expressed polar flagella [31]. When cells are plated on a 1.25% agar surface, lateral flagella are expressed, enabling the cell to swarm on the agar surface. Regulation of lateral flagellar expression is similar to that of biofilm formation and c-di-GMP signaling is involved in the regulation [51] (Supplementary Fig. S7). Gene expression during gelatin-stimulated swarming is thought to be very complex process, involving the c-di-GMP regulation network. So far there have been no insights into how gelatin induces the swarming motility except for the finding that gelatin is involved in the VarS/VarA two-component regulatory system, which regulates the expression of collagenases that degrade gelatin [52]. The homologous GacS/GacA system in *P. aeruginosa* has been shown to be involved in the regulation of flagellar formation and biofilm formation [53]. We speculate that the VarS/VarA system is involved in the production and/or secretion of viscous materials or biosurfactants. *V. alginolyticus* and its relatives are important marine decomposers and are involved in chitin degradation. We showed here that chitin oligosaccharides also stimulated swarming (Supplementary Fig. S1). From the evidence, it is possible that the induction of the degradation systems and the production of swarm-promoting substances, possibly surfactants or viscous polysaccharides might use the same regulation pathway. It is noteworthy that the function of ^laf^FliL, the defect of which prevents swarming but not swimming in agar. The swimming of the ^laf^*fliL* mutant cells in liquid medium was inhibited compared to the wild-type cells and was promoted by the addition of PVP which increases the surrounding viscosity of the cells. The *fliL* protein may affect the gene expression for swarming as shown above (Supplementary Fig. S7). The supplement effects to promote the swarming seem to be too complicated. However, we hope to identify the swarming-promoting factors and how the environmental signals are sensed and the production is regulated in the future.

## MATERIALS AND METHODS

### Strains, plasmids, and media

The bacterial strains and plasmids used here are listed in Table S1. *V. alginolyticus* cells were cultured at 30°C in VC medium (0.5% polypeptone, 0.5% yeast extract, 0.4% K_2_HPO_4_, 3% NaCl, 0.2% glucose), or VPG medium (1% polypeptone, 0.4% K_2_HPO_4_, 3% NaCl, 0.5% glycerol), VNG medium (1% NZ amine, 0.4% K_2_HPO_4_, 3% NaCl, 0.5% glycerol). When necessary, chloramphenicol was added to final concentrations of 2.5 µg/ml for *V. alginolyticus*, after media had been autoclaved. Bacterial cells were suspended in 20TN200 (20mM Tris-HCl, pH8.0, containing 200 mM NaCl).

### Expansion assay of cells by agar plate

VPG or VNG-0.3soft-agar (0.3% agar), VPG or VNG-1.25hard agar (1.25% agar), or VPG or VNG-2.5hard-agar (2.5% agar) agar plate was used for *V. alginolyticus* expansion assay in agar plates. When necessary, Gelatin from bovine bone (Fujifilm, Japan), Casamino acids (BD, USA), Collagen peptides (Nestle, Japan), or Fish collagen peptides TFF-01 (Nippi, Japan) and polyvinylpyrrolidone PVP K90 (TCI, Japan) was added to final concentration of 1% and 5%, respectively and autoclaved. An aliquot (2 µl) of overnight culture was spotted onto a plate, which was then incubated at 30°C in the wetted container. The size of ring expansion was recorded by a scanner. The expansion ring of cells was observed by stereomicroscope (VHX2000, Keyence Co., Osaka, Japan) equipped with high magnification zoom lens (VH-Z450, Keyence Co.) at 3,000-fold magnification. Movies were recorded on a computer (Mac mini) using USB video capture. Those movies are included in the supplementary materials. Only representative data were shown in the Figure 1 and 2, and data reproducibility was confirmed by multiple experiments using biological replicates.

### Observation of swimming behavior

*V. alginolyticus* cells were cultured overnight in VC medium at 30. The overnight culture was suspended in VPG medium at 50- or 100-fold dilution and incubated at 30 for 6h, and cells were observed under a dark-field microscope and recorded by CCD camera on computer or by a digital camera. Swimming profiles were analyzed using a motion analysis software (DIPP-Motion V, DITECT, Japan).

### Transmission electron microscopy

The cells of *V. alginolyticus* were picked from on the agar plates and mixed with 2 % (w/v) potassium phosphotungstate solution (pH 7). The solution was placed on the carbon-coated copper grid and observed with an electron microscope (JEM-1400, JEOL, Japan).

### SDS-PAGE analysis of the cell surface materials

Overnight cultures of YM19 and YM19Δ*fliL* were inoculated onto 1.25% agar-VNG plates with or without 1% gelatin, and incubated at 30°C. After 6 hours, the cells were scraped and suspend in 1 mL of 20TN200. Cell suspensions were stirred using a Polytron for 30 second twice, and centrifuged at 10,000 rpm for 15 min. The supernatant was collected as the samples and was separated by SDS-PAGE, stained with Coomassie Brilliant Blue or and silver, and photographed.

## AUTHOR CONTRIBUTIONS

T.M., S.K., and M.H. designed research; H.N., and M.H. performed experiments; T.M., H.N., S.K., and M.H. analyzed data; S.K., and M.H. wrote the paper.

## DATA AVAILABILITY

The data that supports the findings of this study are available in the Supporting information of this article.

## ACKNOWLEDGEMENTS

We thank Dr. Kimika Maki for technical support with electron microscopy. We thank Dr. David Yadin for English editing of the manuscript. This work was partially supported by JSPS KAKENHI Grant Numbers 23K27140 (to S.K.), and 20H03220 (to M.H.).

## REFERENCES

1. Williams FD, Schwarzhoff RH. 1978. Nature of the swarming phenomenon in *Proteus*. Annu Rev Microbiol 32: 101–122.

2. Fraser GM, Hughes C. 1999. Swarming motility. Curr Opin Microbiol 2: 630–635.

3. Harshey RM. 2003. Bacterial motility on a surface: many ways to a common goal. Annu Rev Microbiol 57: 249–273.

4. Harshey RM, Partridge JD. 2015. Shelter in a swarm. J Mol Biol 427: 3683–3694.

5. Craig L, Forest KT, Maier B. 2019. Type IV pili: dynamics, biophysics and functional consequences. Nat Rev Microbiol 17: 429–440.

6. Ellison CK, Whitfield GB, Brun YV. 2022. Type IV Pili: dynamic bacterial nanomachines. FEMS Microbiol Rev 46: fuab053.

7. McBride MJ. 2019. Bacteroidetes gliding motility and the type IX secretion system. Microbiol Spectr 7: psib-0002-2018.

8. Gorasia DG, Veith PD, Reynolds EC. 2020. The type IX secretion system: advances in structure, function and organisation. Microorganisms 8: 1173.

9. Hölscher T, Kovács ÁT. 2017. Sliding on the surface: bacterial spreading without an active motor. Environ Microbiol 19: 2537–2545.

10. Kearns DB. 2010. A field guide to bacterial swarming motility. Nat Rev Microbiol 8: 634–644.

11. Partridge JD, Harshey RM. 2013. Swarming: flexible roaming plans. J Bacteriol 195: 909–918.

12. Jaques S, McCarter LL. 2006. Three new regulators of swarming in *Vibrio parahaemolyticus*. J Bacteriol 188: 2625–2635.

13. Ulitzur S. 1974. Induction of swarming in *Vibrio parahaemolyticus*. Arch Microbiol 101: 357–363.

14. Kearns DB, Losick R. 2003. Swarming motility in undomesticated *Bacillus subtilis*. Mol Microbiol 49: 581–590.

15. Harshey RM, Matsuyama T. 1994. Dimorphic transition in *Escherichia coli* and *Salmonella typhimurium*: surface-induced differentiation into hyperflagellate swarmer cells. Proc Natl Acad Sci USA 91: 8631–8635.

16. Deditius JA, Felgner S, Spöring I, Kühne C, Frahm M, Rohde M, Weiß S, Erhardt M. 2015. Characterization of novel factors involved in swimming and swarming motility in *Salmonella enterica* Serovar Typhimurium. PLoS One 10: e0135351.

17. Be’er A, and Ariel G. 2019. A statistical physics view of swarming bacteria. Mov Ecol 7: 9.

18. Bhagwat AA, Young L, Smith AD, Bhagwat M. 2017. Transcriptomic analysis of the swarm motility phenotype of *Salmonella enterica* Serovar Typhimurium mutant defective in periplasmic glucan synthesis. Curr Microbiol 74: 1005–1014.

19. Little K, Austerman J, Zheng J, Gibbs KA. 2019. Cell shape and population migration are distinct steps of proteus mirabilis swarming that are decoupled on high-percentage agar. J Bacteriol 201: e00726–18.

20. Wu Y, Berg HC. 2012. Water reservoir maintained by cell growth fuels the spreading of a bacterial swarm. Proc Natl Acad Sci USA 109: 4128–4133.

21. Ping L, Wu Y, Hosu BG, Tang JX, Berg HC. 2014. Osmotic pressure in a bacterial swarm. Biophys J 107: 871–878.

22. Minamino T, Kinoshita M. 2023. Structure, assembly, and function of flagella responsible for bacterial locomotion. EcoSal Plus 11: eesp00112023.

23. Homma M, Kojima S. 2022. Roles of the second messenger c-di-GMP in bacteria: Focusing on the topics of flagellar regulation and *Vibrio* spp. Genes Cells 27: 157–172.

24. Harshey RM, Toguchi A. 1996. Spinning tails: homologies among bacterial flagellar systems. Trends Microbiol 4: 226–231.

25. Minamino T, Kinoshita M, Namba K. 2019. Directional switching mechanism of the bacterial flagellar motor. Comput Struct Biotechnol J 17: 1075–1081.

26. Armitage JP, Berry RM. 2020. Assembly and dynamics of the bacterial flagellum. Annu Rev Microbiol 74: 181–200.

27. Terashima H, Kojima S, Homma M. 2008. Flagellar motility in bacteria structure and function of flagellar motor. Int Rev Cell Mol Biol 270: 39–85.

28. Takekawa N, Imada K, Homma M. 2020. Structure and energy-conversion mechanism of the bacterial Na^+^-driven flagellar motor. Trends Microbiol 28: 719–731.

29. Kojima S, Shinohara A, Terashima H, Yakushi T, Sakuma M, Homma M, Namba K, Imada K. 2008. Insights into the stator assembly of the *Vibrio* flagellar motor from the crystal structure of MotY. Proc Natl Acad Sci USA 105: 7696–7701.

30. Atsumi T, McCarter L, Imae Y. 1992. Polar and lateral flagellar motors of marine *Vibrio* are driven by different ion-motive forces. Nature 355: 182–184.

31 Kawagishi I, Maekawa Y, Atsumi T, Homma M, Imae Y. 1995. Isolation of the polar and lateral flagellum-defective mutants in *Vibrio alginolyticus* and identification of their flagellar driving energy sources. J Bacteriol 177: 5158–5160.

32. Atsumi T, Maekawa Y, Yamada T, Kawagishi I, Imae Y, Homma M. 1996. Effect of viscosity on swimming by the lateral and polar flagella of *Vibrio alginolyticus*. J Bacteriol 178: 5024–5026.

33. Kawagishi I, Imagawa M, Imae Y, McCarter L, Homma M. 1996. The sodium-driven polar flagellar motor of marine *Vibrio* as the mechanosensor that regulates lateral flagellar expression. Mol Microbiol 20: 693–699.

34. McCarter L, Hilmen M, Silverman M. 1988. Flagellar dynamometer controls swarmer cell differentiation of *V. parahaemolyticus*. Cell 54: 345–351.

35. Merino S, Shaw JG, Tomás JM. 2006. Bacterial lateral flagella: an inducible flagella system. FEMS Microbiol Lett 263: 127–135.

36. De Boer WE, Golten C, Scheffers WA. 1975. Effects of some physical factors on flagellation and swarming of *Vibrio alginolyticus*. Netherlands Journal of Sea Research 9: 197–213.

37. Partridge JD, Harshey RM. 2024. Flagellar protein FliL: A many-splendored thing. Mol Microbiol 122: 447–454.

38. Takekawa N, Isumi M, Terashima H, Zhu S, Nishino Y, Sakuma M, Kojima S, Homma M, Imada K. 2019. Structure of *Vibrio* FliL, a new stomatin-like protein that assists the bacterial flagellar motor function. mBio 10: e00292–00219.

39. Zhu S, Kumar A, Kojima S, Homma M. 2015. FliL associates with the stator to support torque generation of the sodium-driven polar flagellar motor of *Vibrio*. Mol Microbiol 98: 101–110.

40. Lin TS, Zhu S, Kojima S, Homma M, Lo CJ. 2018. FliL association with flagellar stator in the sodium-driven *Vibrio* motor characterized by the fluorescent microscopy. Sci Rep 8: 11172.

41. Daumke O, Lewin GR. 2022. SPFH protein cage - one ring to rule them all. Cell Res 32: 117–118.

42. Yokoyama H, Matsui I. 2023. Higher-order structure formation using refined monomer structures of lipid raft markers, Stomatin, Prohibitin, Flotillin, and HflK/C-related proteins. FEBS Open Bio 13: 926–937.

43. Lama H, Yamamoto MJ, Furuta Y, Shimaya T, Takeuchi KA. 2024. Emergence of bacterial glass. PNAS Nexus 3: pgae238.

44. Aranson IS. 2022. Bacterial active matter. Rep Prog Phys 85: 076601.

45. Wang Y, Bian Z, Wang Y. 2022. Biofilm formation and inhibition mediated by bacterial quorum sensing. Appl Microbiol Biotechnol 106: 6365–6381.

46. Fung BL, Esin JJ, Visick KL. 2024. *Vibrio fischeri*: a model for host-associated biofilm formation. J Bacteriol 206: e0037023.

47. Niu C, Graves JD, Mokuolu FO, Gilbert SE, Gilbert ES. 2005. Enhanced swarming of bacteria on agar plates containing the surfactant Tween 80. J Microbiol Methods 62: 129–132.

48. Mignot T, Shaevitz JW, Hartzell PL, Zusman DR. 2007. Evidence that focal adhesion complexes power bacterial gliding motility. Science 315: 853–856.

49. Agrebi R, Wartel M, Brochier-Armanet C, Mignot T. 2015. An evolutionary link between capsular biogenesis and surface motility in bacteria. Nat Rev Microbiol 13: 318–326.

50. McCarter L, Silverman M. 1990. Surface-induced swarmer cell differentiation of V*ibrio parahaemolyticus*. Mol Microbiol 4: 1057–1062.

51. Homma M, Nishikino T, Kojima S. 2022. Achievements in bacterial flagellar research with focus on *Vibrio* species. Microbiol Immunol 66: 75–95.

52. Mima T, Gotoh K, Yamamoto Y, Maeda K, Shirakawa T, Matsui S, Murata Y, Koide T., Tokumitsu H, Matsushita O. 2018. Expression of collagenase is regulated by the VarS/VarA two-component regulatory system in *Vibrio alginolyticus*. J Membr Biol 251: 51–63.

53. Song H, Li Y, Wang Y. 2023. Two-component system GacS/GacA, a global response regulator of bacterial physiological behaviors. Eng Microbiol 3: 100051.

54. Berg HC. 2005. Swarming motility: it better be wet. Curr Biol 15: R599–600.

55. Turner L, Zhang R, Darnton NC, Berg HC. 2010. Visualization of Flagella during bacterial Swarming. J Bacteriol 192: 3259–3267.

56. Böttcher T, Elliott HL, Clardy J. 2016. Dynamics of Snake-like Swarming Behavior of *Vibrio alginolyticus*. Biophys J 110: 981–992.

