## Supplemental Figure 1 for "Requirements for swarming ability by lateral flagella on an agar surface in marine *Vibrio* cells"

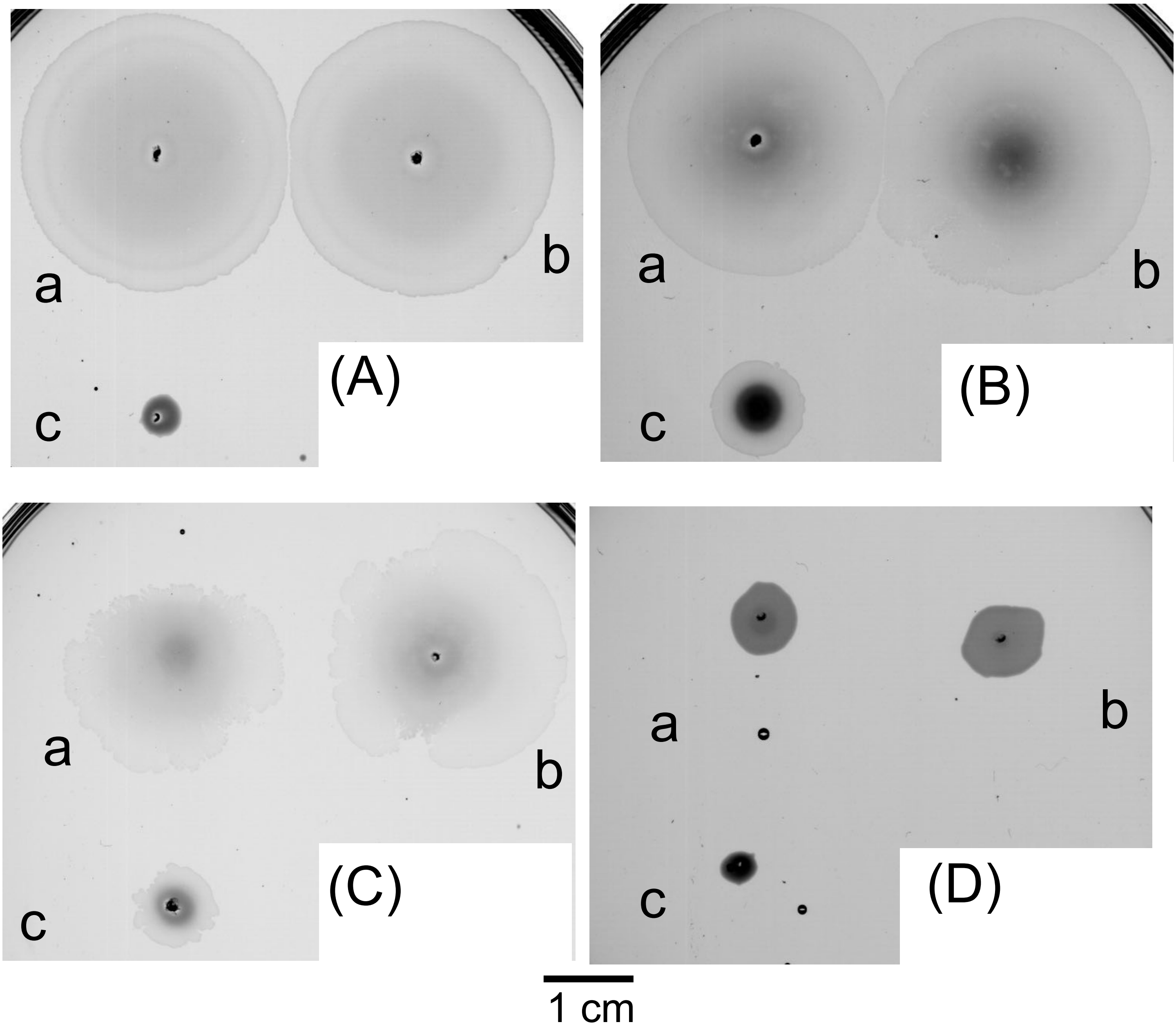


**Fig. S1.** Swarming profiles of *Vibrio* cells on 1.25% agar containing various supplements. 2 μL of the overnight culture (a:138-2, b:YM19, c:YM19$^{laf}$*fliL*) was put on the 1.25% agar-VNG plate added 200μL of 10% chitin oligosaccharides (A), 200μL of 10% beef collagen peptides (B), 200μL of 10% fish collagen peptides (C) or none (D), and incubated at 30° C for 5 hours. The plates were scanned with a photo scanner.
