## Supplemental Figure 2 for "Requirements for swarming ability by lateral flagella on an agar surface in marine *Vibrio* cells"

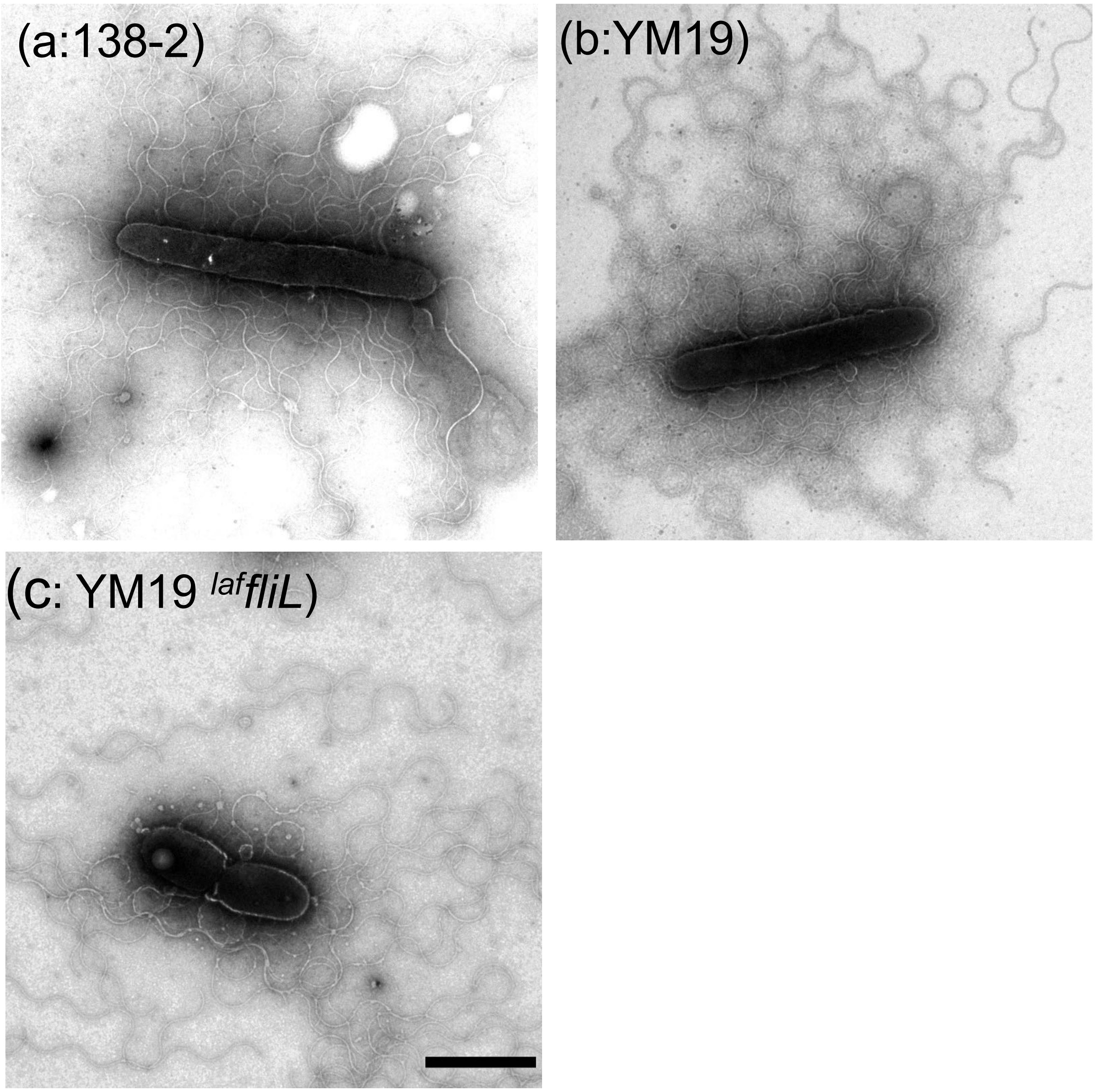


**Fig. S2.** Flagella formation observed by electron microscopy. 2 $\mu$L of the overnight culture of 138-2 (a), YM19 (b), or YM19$^{laf}$*fliL* (c) was put on the 1.25% agar-VPG plate and incubated at 30° C overnight. The edges of the colonies were picked up with a toothpick, suspended in 0.5% PTA, and observed under an electron microscope. Bar shows 2$\mu$m.
