## Supplemental Figure 3 for "Requirements for swarming ability by lateral flagella on an agar surface in marine *Vibrio* cells"

### Slide 1
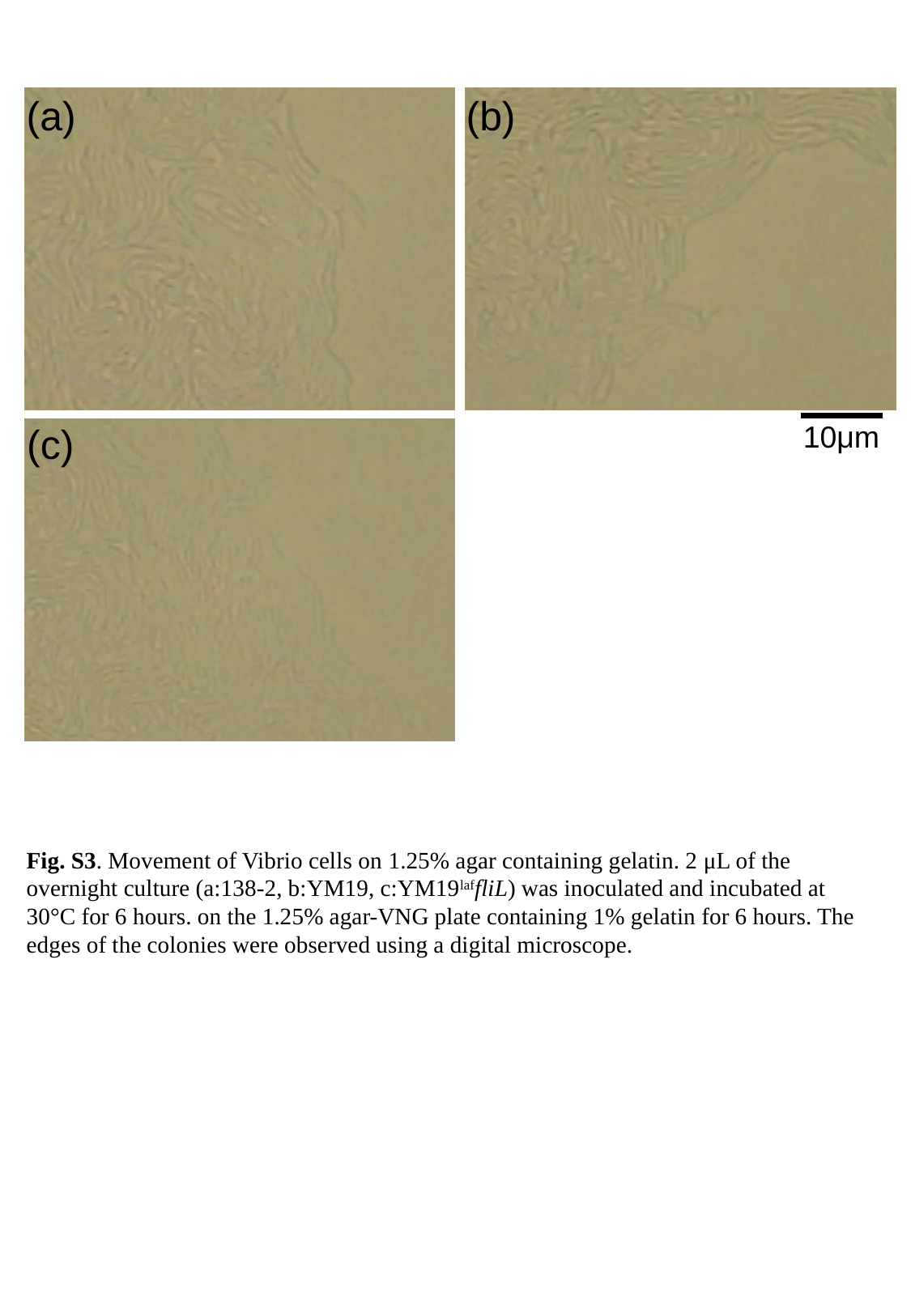

(b)
(a)
10μm
(c)
Fig. S3. Movement of Vibrio cells on 1.25% agar containing gelatin. 2 μL of the overnight culture (a:138-2, b:YM19, c:YM19^laf^fliL) was inoculated and incubated at 30°C for 6 hours. on the 1.25% agar-VNG plate containing 1% gelatin for 6 hours. The edges of the colonies were observed using a digital microscope.
