## Supplemental Figure 4 for "Requirements for swarming ability by lateral flagella on an agar surface in marine *Vibrio* cells"

### Slide 1
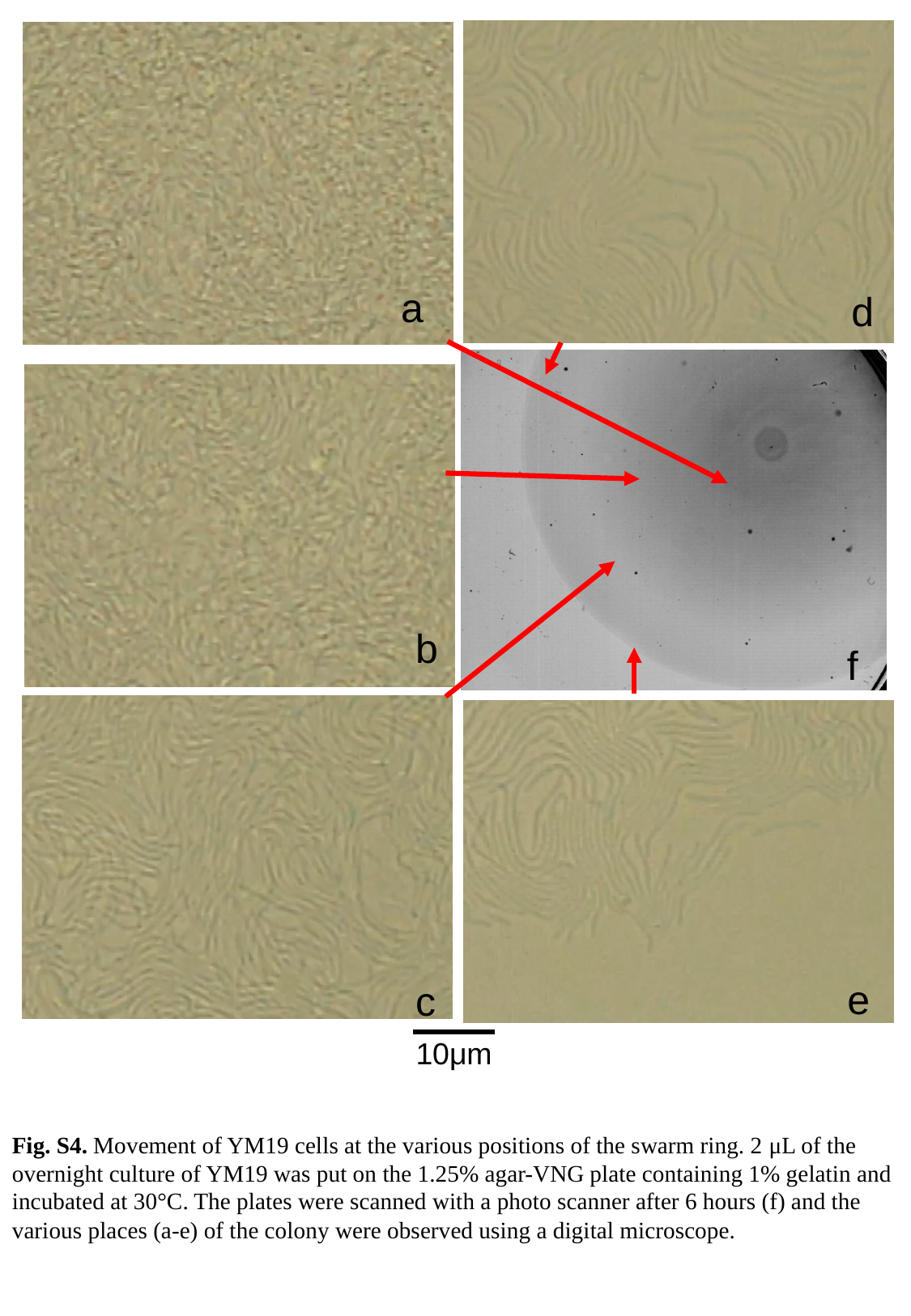

a
d
b
f
e
c
10μm
Fig. S4. Movement of YM19 cells at the various positions of the swarm ring. 2 μL of the overnight culture of YM19 was put on the 1.25% agar-VNG plate containing 1% gelatin and incubated at 30°C. The plates were scanned with a photo scanner after 6 hours (f) and the various places (a-e) of the colony were observed using a digital microscope.
