## Supplemental Figure 5 for "Requirements for swarming ability by lateral flagella on an agar surface in marine *Vibrio* cells"

### Slide 1
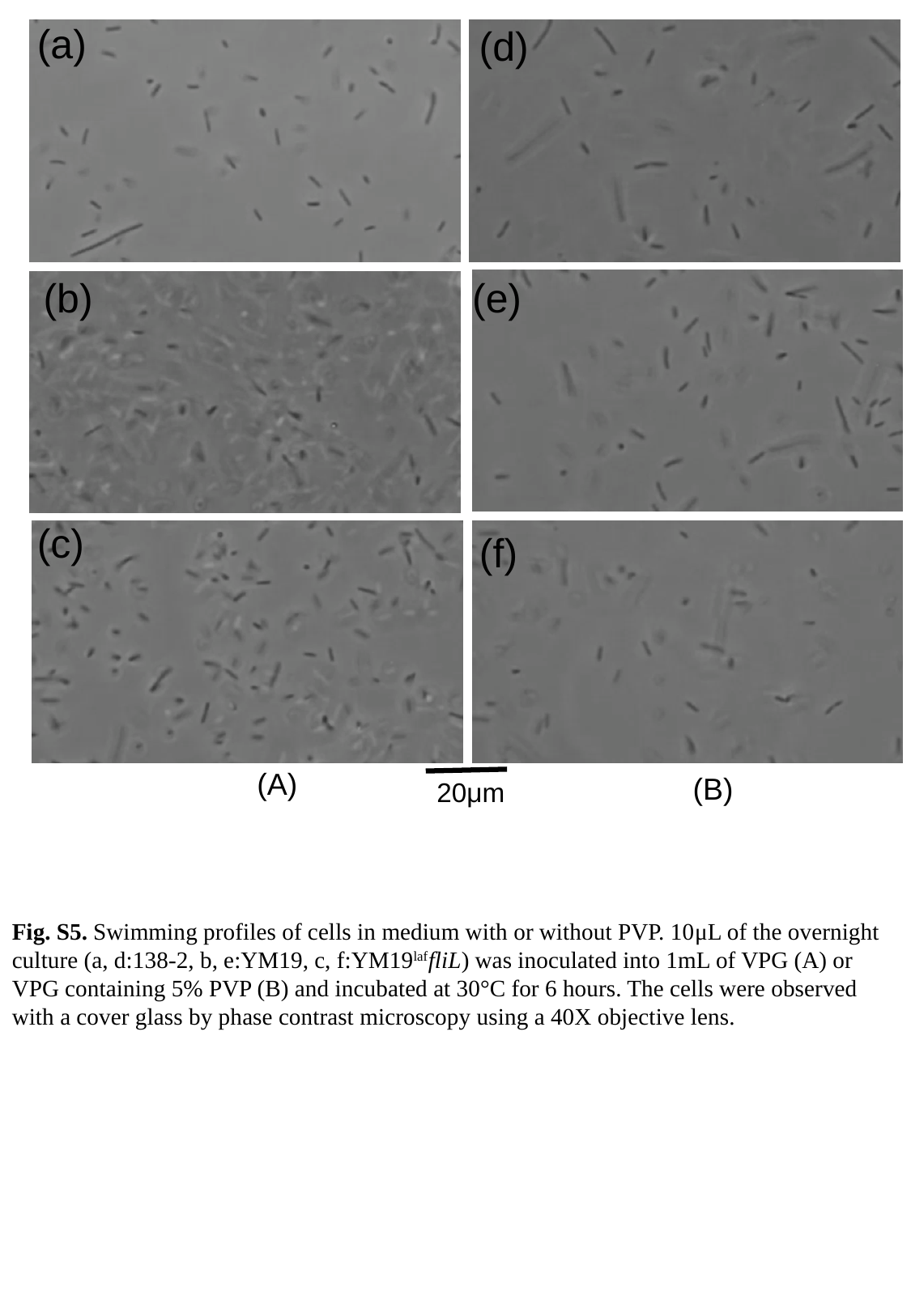

(a)
(d)
(e)
(b)
(c)
(f)
(A)
(B)
20μm
Fig. S5. Swimming profiles of cells in medium with or without PVP. 10μL of the overnight culture (a, d:138-2, b, e:YM19, c, f:YM19^laf^fliL) was inoculated into 1mL of VPG (A) or VPG containing 5% PVP (B) and incubated at 30°C for 6 hours. The cells were observed with a cover glass by phase contrast microscopy using a 40X objective lens.
