## Supplemental Figure 6 for "Requirements for swarming ability by lateral flagella on an agar surface in marine *Vibrio* cells"

### Slide 1
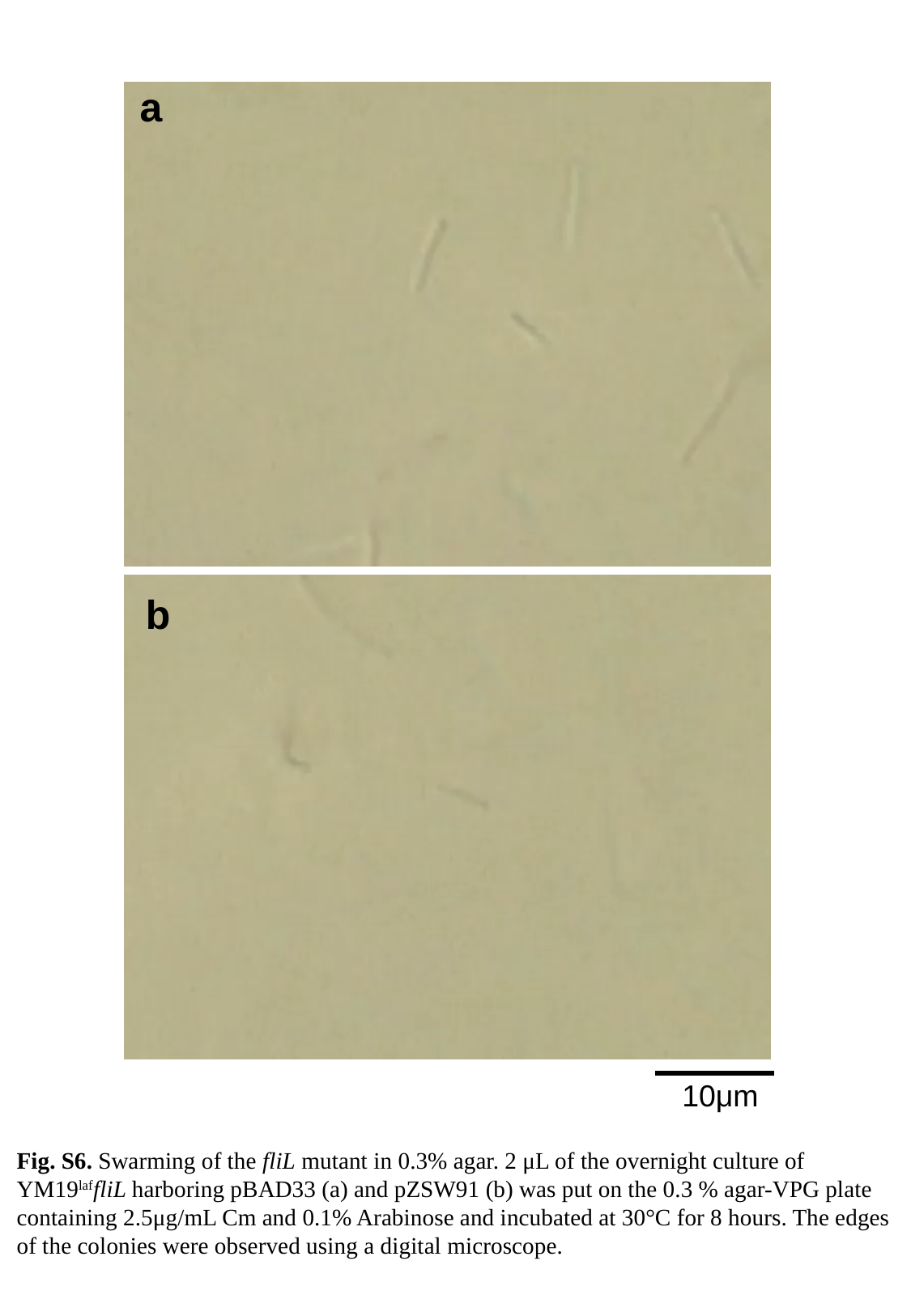

a
b
10μm
Fig. S6. Swarming of the fliL mutant in 0.3% agar. 2 μL of the overnight culture of YM19^laf^fliL harboring pBAD33 (a) and pZSW91 (b) was put on the 0.3 % agar-VPG plate containing 2.5μg/mL Cm and 0.1% Arabinose and incubated at 30°C for 8 hours. The edges of the colonies were observed using a digital microscope.
