## Supplemental Figure 7 for "Requirements for swarming ability by lateral flagella on an agar surface in marine *Vibrio* cells"

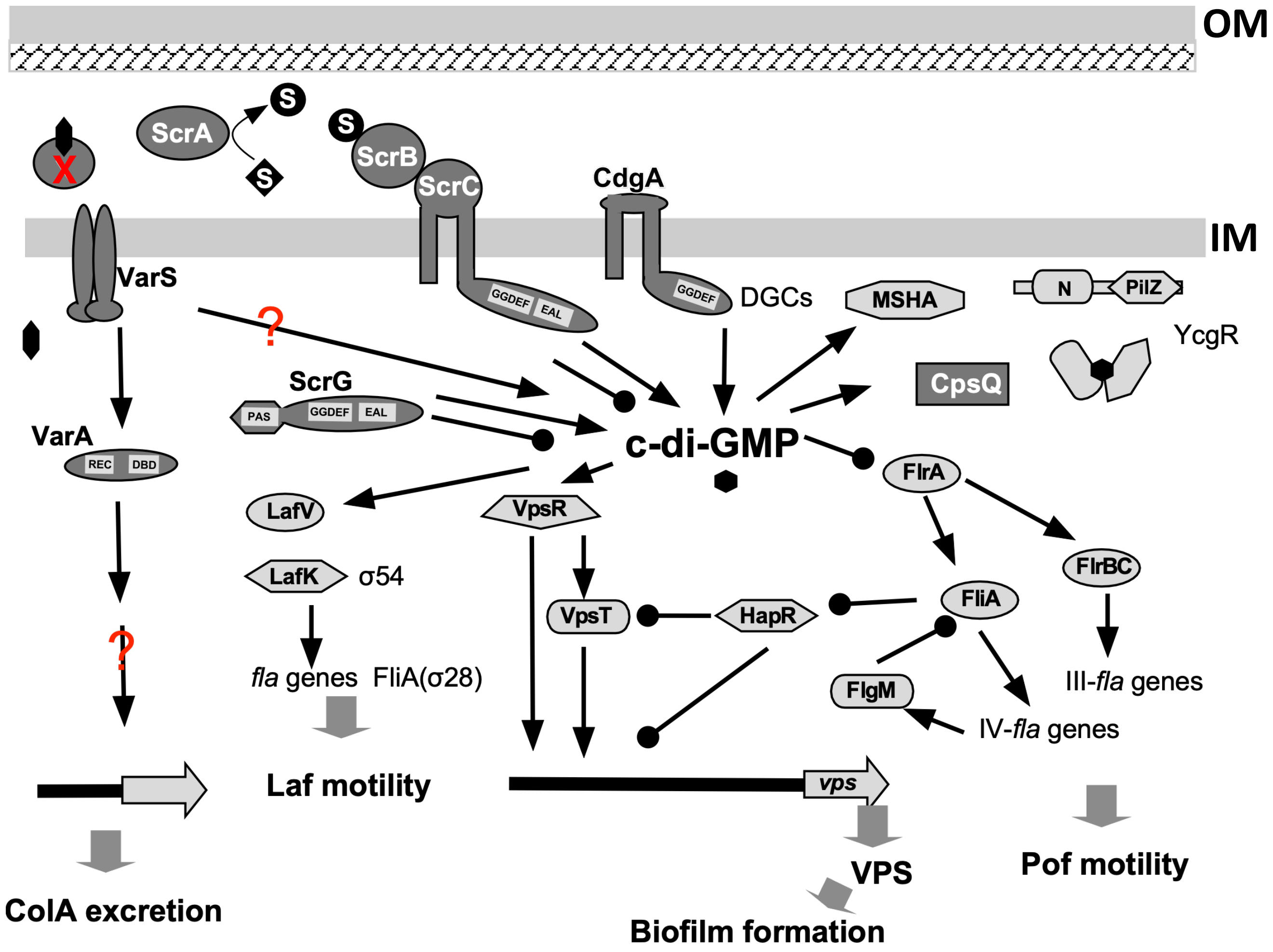


**Fig. S7.** Possible regulation of lateral flagella, polar flagella, biofilm, or collagen in *Vibrio* cells. The arrow ends represent positive regulation, and the black circle ends represent negative regulation. IM; inner membrane, OM; outer membrane, VPS; *Vibrio* polysaccharide, Pof; polar flagella, Laf; lateral flagella, X (in red); unknown protein for collagen or collagen peptides.
